# Evolution of protease activation and specificity via alpha-2-macroglobulin-mediated covalent capture

**DOI:** 10.1101/2023.01.19.524706

**Authors:** Philipp Knyphausen, Mariana Rangel-Pereira, Paul Brear, Marko Hyvönen, Lutz Jermutus, Florian Hollfelder

## Abstract

Tailoring of the activity and specificity of proteases is critical for their utility across industrial, medical and research purposes. However, engineering or evolving protease catalysts is challenging and often labour intensive. Here, we describe a generic method to accelerate this process based on yeast display. We introduce the protease selection system A2M^cap^ that covalently captures protease catalysts by repurposed alpha-2-macroglobulin (A2Ms). To demonstrate the utility of A2M^cap^ for protease engineering we exemplify the directed activity and specificity evolution of six serine proteases. This resulted in a variant of *Staphylococcus aureus* serin-protease-like (Spl) protease SplB, an enzyme used for recombinant protein processing that no longer requires activation by N-terminal signal peptide removal. SCHEMA-based domain shuffling was used to map the specificity determining regions of Spl proteases leading to a chimeric scaffold that supports specificity switching via subdomain exchange. The ability of A2M^cap^ to overcome key challenges *en route* to tailor-made proteases suggests easier access to such reagents in the future.

## Introduction

Proteases are proteolytic enzymes that perform essential regulatory and metabolic functions in all domains of life. As a consequence, proteases find application as therapeutic agents^1^, as targets or constituents of activatable biologics^2^, but also as tools for biomedical research, biotechnological applications and synthetic biology^3,4^. However, only a minority of proteases are available for each of these applications, often displaying suboptimal properties in terms of stability, activity, specificity or substrate scope. There is thus a need for efficient protein engineering methods that enable improvements of existing proteases and their reprogramming towards novel substrates.

Protein binders for a range of therapeutic applications can be easily generated by ‘panning’, in which protein libraries are allowed to bind to a target antigen and non-binders subsequently removed by washing. Multiple high-throughput platforms are suitable for this selection process, e.g. phage-, ribosome- or yeast surface display and have become workhorses for the routine discovery of affinity reagents. No similarly general protein display technology exists that can select for target binding *and* cleavage, and of the currently existing systems for protease directed evolution, none fully leverages the flexibility and versatility of panning-based selection. Instead, many current systems are based on co-expression of the protease and a readout substrate^5–9^ or on non-generalizable suicide inhibitors as substrates^10^. Such setups limit control over factors that define selection pressure, e.g. substrate concentration and incubation time.

Here we describe the use of bacterial alpha-2-macroglobulins (A2Ms) as generic covalent capture probes to identify protease variants in an analogous way to panning-based selection for binders. As part of metazoan defense systems against invasive bacteria, A2Ms, which are large proteins of >150 kDa, trap any protease covalently upon cleavage of a peptide bait sequence in a mechanism reminiscent of Venus flytraps. Bait cleavage results in a conformational change and concomitant exposure of a reactive thioester, which in turn attacks primary amines on the surface of the incoming protease, leading to the formation of a covalent protease-inhibitor complex^11–13^. As their metazoan counterparts, bacterial A2Ms act as covalent protease inhibitors but are monomeric and more amenable to recombinant expression and bait sequence manipulation^14–17^. We take advantage of these properties by developing A2M-based protease capture (A2M^cap^) as a tool for characterisation and broad spectrum directed evolution of proteases. To showcase A2M^cap^, we evolved functional chimeras of the six closely related serine-protease-like proteases SplA-F of *Staphylococcus aureus*^18,19^ and mapped their specificity-determining subdomain. In a separate directed evolution campaign, we selected SplB mutants that do not require post-translational pre-form activation at the N-terminus. These mutants enable direct expression of active enzyme and thus pave the way for a simplified use of SplB in biotechnological and synthetic biology applications.

## Results

### A2M^cap^ measures protease activity by covalent substrate trapping

The principle behind A2M^cap^ is the formation of a covalent bond between a protease of interest and a modified A2M substrate. A covalent bond is formed if a protease cleaves the A2M bait sequence, which triggers a structural rearrangement in A2M and subsequent reaction of its Cys-Gln thioester with surface lysines of the protease, thereby trapping it covalently. We chose *Escherichia coli* A2M, in which a part of the naturally present bait sequence is replaced with a protease recognition site of choice. Additional modifications on A2M include appropriate tags for purification (10x-His-tag) and labelling (Avi-tag for *in vivo* biotinylation) (Supplementary Fig. 1a). After recombinant production in *E*.*coli*, these modified A2Ms were used as substrates. Proteases of interest were displayed on the surface of yeast cells, where A2M capture through bait-cleavage and the protease display levels are detectable with fluorescent labels. This setup enables measurements of catalytic turnover via flow cytometry (Fig. 1c; exemplary gating strategy shown in Supplementary Fig. 1b).

**Figure 1:**
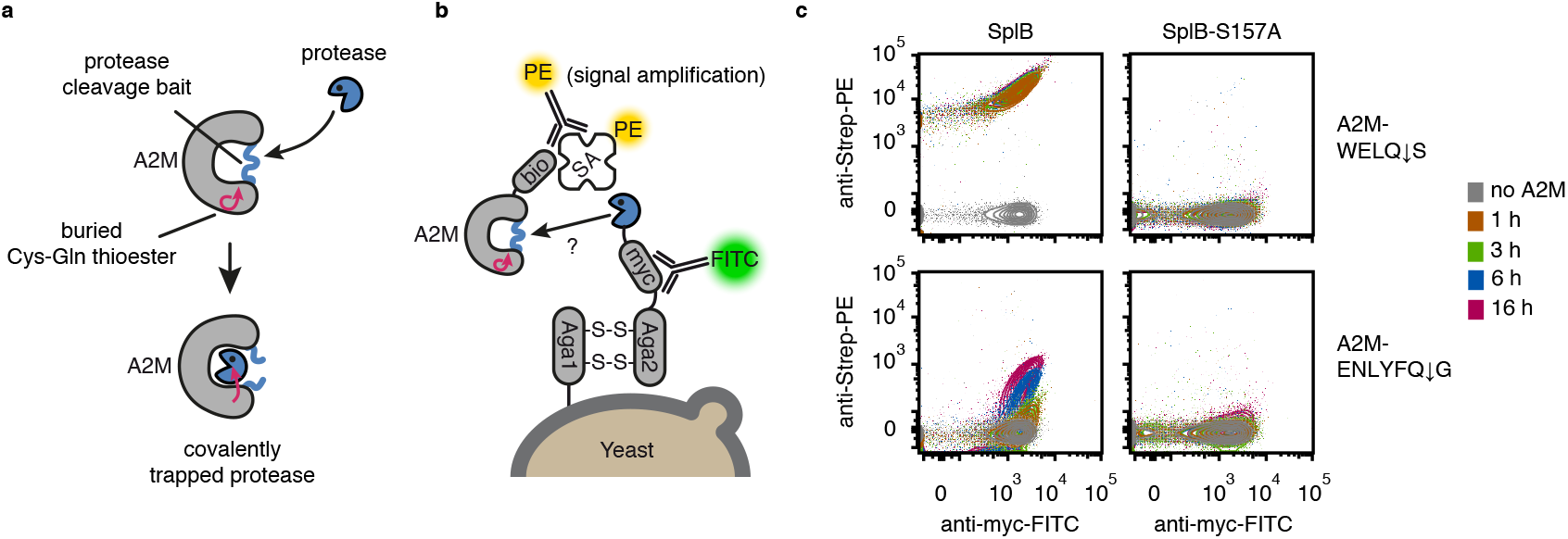
A2M^cap^ enables protease activity detection on the yeast cell surface. **(a)** Principle of alpha-2-macroglobulin (A2M)-based protease trapping. Cleavage of the bait sequence leads to a structural rearrangement in A2M that enables reaction of cysteine-glutamine (Cys-Gln) thioester with surface lysines of the protease, thereby trapping it covalently. The bait sequence (depicted as a blue line) can be replaced by a protease cleavage site of choice. **(b)** Setup of turnover selection for libraries of yeast-displayed protease. Proteases (blue pacman) are displayed on the surface of yeast cells via an Aga2 fusion (which attaches to the yeast cell surface via disulfide bonds to Aga1; indicated by S-S); optical label are incorporated for simultaneous quantification of protease display level and catalytic turnover, together providing a measurement of normalised activity: (i) The level of protease display is read-out via a fluorescein isothiocyanate (FITC)-conjugated anti-myc tag antibody (Y-shaped antibody symbol with green label); (ii) Protease activity is probed by incubating yeast cells with biotin-tagged A2M. The captured A2M molecules are subsequently labelled with Phycoerythrin (PE)-streptavidin (SA), which binds the biotin moiety. Optionally, the signal may be amplified through a PE-anti-streptavidin antibody. **(c)** A2M^cap^-based detection of protease activity via flow cytometry. SplB-displaying yeast cells were reacted with A2M-WELQ↓S or A2M-ENLYFQ↓S (cognate recognition sequence of TEV protease, ENLYFQ↓S, serves as negative control; low activity of SplB expected towards this substrate). SplB-S157A is a mutant that is displayed on yeast, but inactive as a protease (n=1 flow cytometry measurement). Source data are provided as a Source Data file for panel c.

As a target protease we chose the WELQ↓X-specific serine protease SplB from *S. aureus*^20^, which was displayed and probed with A2M bearing a WELQ↓S bait (A2M-WELQ↓S) or a control bait (A2M-ENLYFQ↓S, i.e. an A2M with a TEV cleavage site instead). With the cognate substrate A2M-WELQ↓S, we observed an A2M-specific signal increase over two orders of magnitude within 10 min and reaching saturation after 1 h. By contrast, even after 16 h of incubation with A2M-ENLYFQ↓S the signal remained about 30-fold lower. Yeast cells displaying an inactive SplB mutant (S157A) did not show a signal with either substrate (Fig. 1c and Supplementary Fig. 1c). The observation of time-resolved readouts for protease activity and specificity suggests that, by adjusting the reaction time, selection stringency can be controlled for directed evolution applications.

### Directed evolution of an SplB variant that does not require N-terminal processing

Like many proteases, SplB is translated in an inactive form and post-translational processing at the N-terminus (to remove its signal peptide and expose N-terminal residue Glu-37) is required to become fully active. In fact, any extension at Glu-37 has an inhibitory effect on SplB, which limits its utility in synthetic biology and other applications^18^. To address and overcome this limitation, we sought to employ A2M^cap^ for the directed evolution of SplB variants that would not require post-translational activation.

We designed an SplB display construct that mimics the immature form (pre-SplB) due to an N-terminal Aga2 fusion that prevents correct processing (Fig. 2a). When probed with A2M-WELQ↓S substrate for 12h, pre-SplB yielded about 100-fold lower product signal than SplB, despite similar levels of protease display (Fig. 2b). The difference between pre-SplB and SplB was also reflected in their enrichment behaviour in magnetic-activated cells sorting experiments (MACS): SplB-displaying cells were readily enriched from a 1000-fold excess of cells displaying inactive SplB S157A (enrichment factor ~300-fold), while no enrichment was apparent for pre-SplB (Fig. 2c). The observed enrichments established A2M^cap^ as a suitable high-throughput selection system for the directed evolution of SplB variants not requiring N-terminal processing.

**Figure 2:**
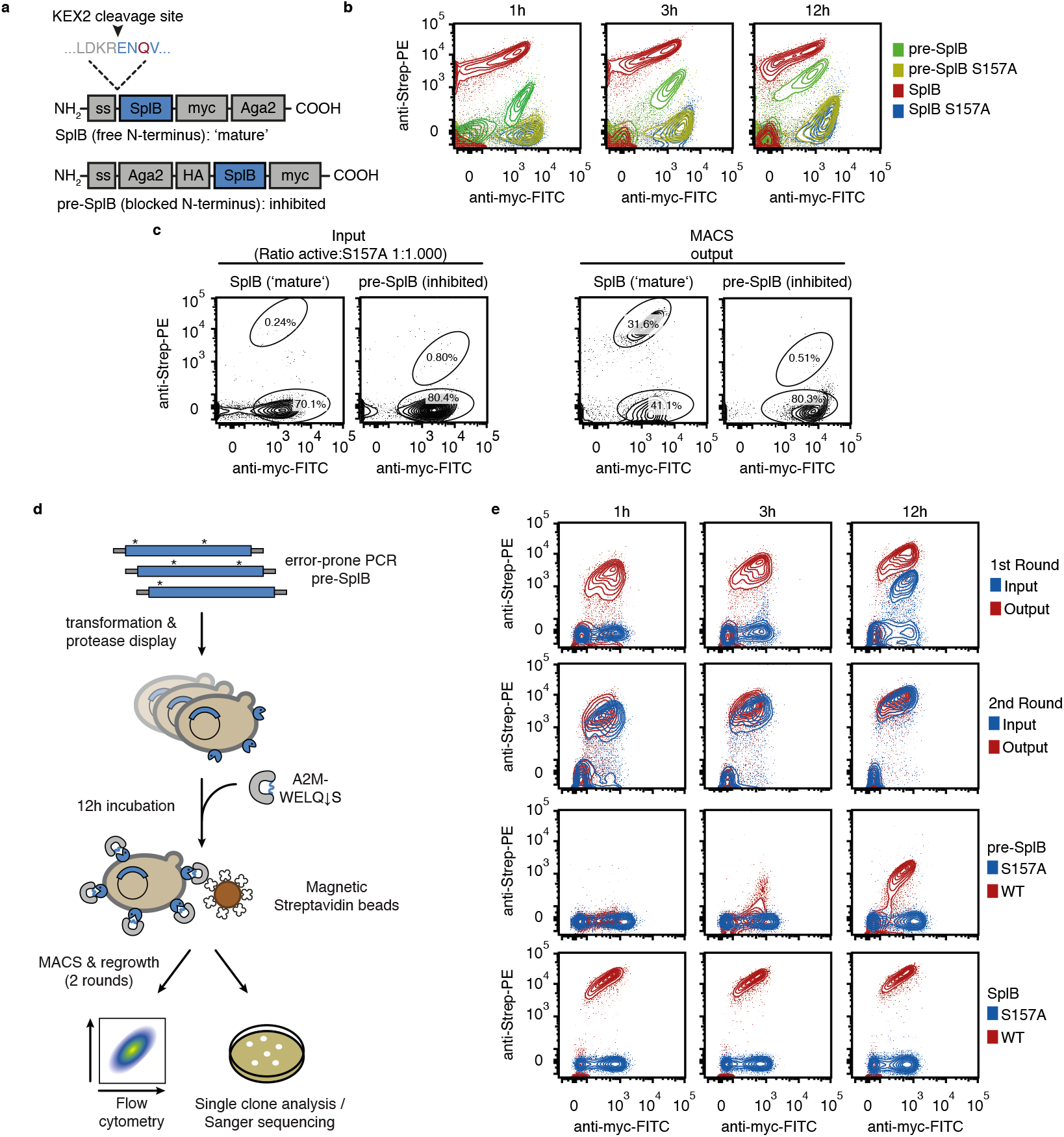
Directed evolution of N-terminal processing-independent SplB using A2M^cap^. **(a)** Schematic depiction of SplB constructs used for yeast surface display. *Top*: ‘Mature’ SplB is expressed with a KEX2 cleavage site directly upstream of the N-terminal Glu of mature SplB and a C-terminal Aga2 fusion as a display anchor (SplB residues in blue, N3Q mutation of a predicted glycosylation site shown in red). *Bottom*: pre-SplB is expressed with an N-terminal Aga2 fusion, blocking the N-terminus and leading to an inhibited enzyme. **(b)** Flow cytometric comparison of ‘mature’ SplB and pre-SplB using A2M-WELQ↓S as a substrate, incubated for 1h, 3h or 12h. Inactive forms of both forms (S157A) are used as negative controls (representative data of *n*=2 independent inductions and flow cytometry measurements). **(c)** Proof-of-concept for selection using magnetic activated cell sorting (MACS) of ‘mature’ SplB and pre-SplB. Left panel: Input populations comprised 1:1000 mixtures of active and inactive (S157A) SplB. Right panel: Output populations after MACS (*n*=1 MACS and flow cytometry measurement). **(d)** Overview of mutagenesis and selection strategy used for directed evolution of pre-SplB. After error-prone PCR, cloning and display, yeast cells were incubated with A2M-WELQ↓S for 12h, washed and selected via MACS. After each round, cells were regrown and analysed by flow cytometry. Isolated clones were analysed after the second round (representative data of *n*=2 independent inductions and flow cytometry measurements). **(e)** Top panel: Flow cytometric analysis of 1^st^ and 2^nd^ selection round. Bottom panel: pre-SplB and ‘mature’ SplB measured under the same conditions as the input and output population in top panel. FITC = Fluorescein isothiocyanate, Strep-PE = Streptavidin-Phycoerythrin. Source data are provided as a Source Data file for panel b and c.

In the absence of a precise mechanism of pre-SplB inhibition, we opted for a random mutagenesis approach using error-prone PCR. A library with ~1.9 mutations/gene and a final diversity of ~2×10^8^ cfus was screened against A2M-WELQ↓S using two rounds of MACS (Fig. 2d). We observed a ~140-fold increase in the overall population activity after the first round of enrichment, followed by a further 1.5-fold improvement in the second round (Fig. 2e). Sequencing of 14 randomly picked single clones revealed that mutations at two sites (E89 and S154) were present in 11 and 4 clones, respectively (Supplementary Fig. 2a). Neither site had previously been implicated in SplB activation^21^. The residue S154 is located close to the active site serine S157 while E89 is located about 25 Å from any SplB active site residue (PDB 2VID, Fig. 3a). Kinetic analysis of the six selected variants in purified form, suggested that the mutation S154R had the strongest impact on activity *in vitro* (Fig. 3b, Supplementary Fig. 2b). The best hit, the double mutant N2K/S154R, showed an improvement of 11-fold over wildtype pre-SplB (while approaching wild-type mature SplB (~26%; *k*_cat_/*K*_m_=1670 and 6550 M^-1^ s^-1^, respectively). Mass spectroscopic analysis suggested that the activity improvement of S154R mutants is not a result of increased spurious N-terminal processing compared to the wild type protein (Supplementary Fig. 2c, Supplementary Table 1). The selection of genuinely improved mutants demonstrates that A2M^cap^ selects proteases with higher catalytic activity.

**Figure 3:**
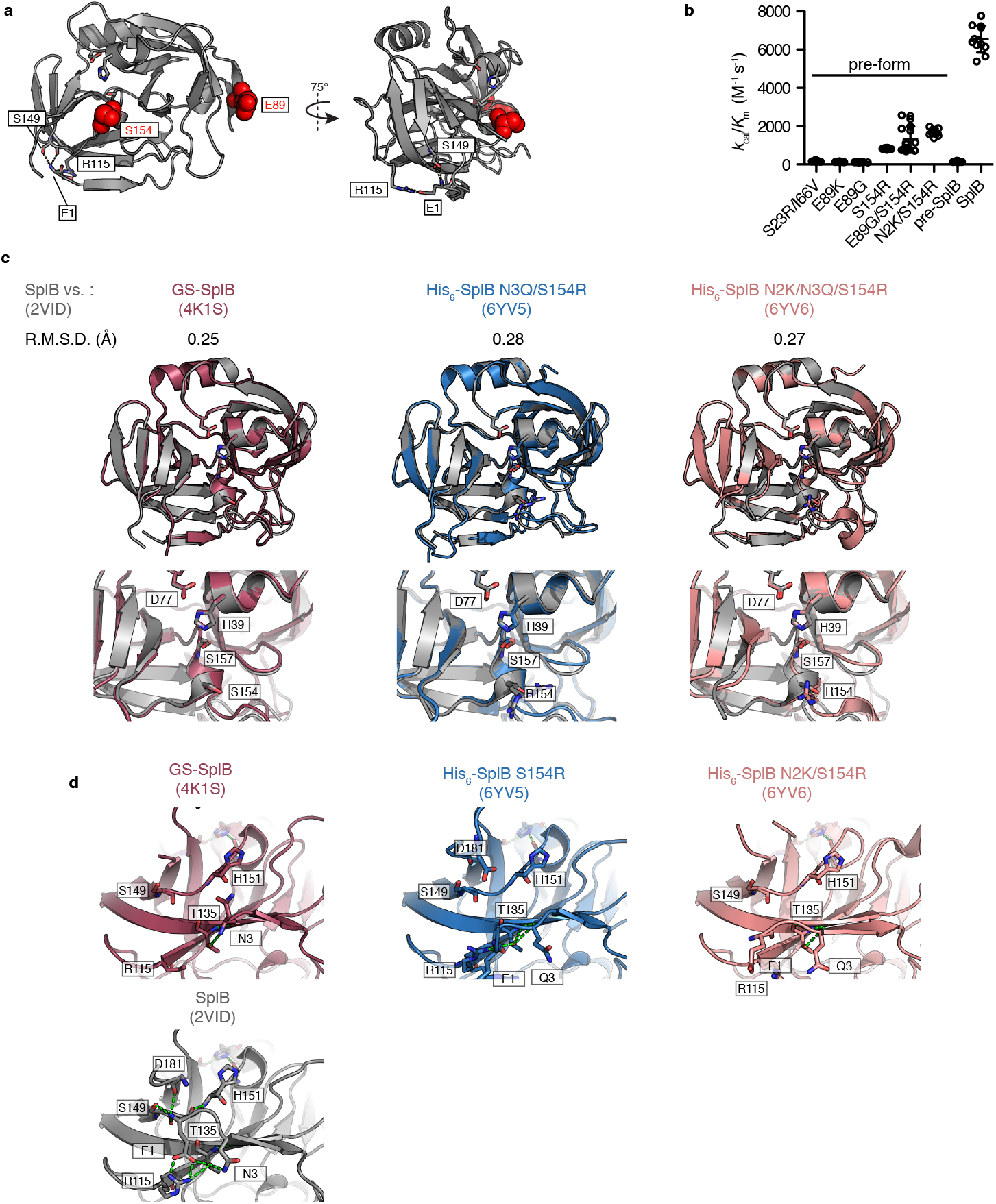
Structure and activity of evolved pre-SplB variants. **(a)** SplB structure (PDB: 2VID) with positions most enriched in directed evolution campaign highlighted in red. Positions previously implicated in SplB activation (black) and the underlying hydrogen bond network are also shown. **(b)** Michaelis-Menten kinetics of pre-SplB mutants and mature SplB (*n*=3 independent measurements of the same protein purification, mean and SD). **(c)** Structural alignment of active SplB (PDB: 2VID) with pre-form SplB variants GS-SplB (PDB: 4K1S), His_6_-SplB-E3Q/S154R (PDB: 6YV5) and His_6_-SplB-E2K/E3Q/S154R (PDB: 6YV6). The root-mean-square deviation (RMSD) is indicated for each alignment. Top panel: Overview of the whole structure. Bottom panel: Close-up of the catalytic triad. **(d)** Detailed view of the hydrogen bonds formed involving the N-terminus. The residues that were described by Pustelny et al.^21^ to be important for SplB activation are highlighted. Hydrogen bonds are indicated as green dashes. Source data are provided as a Source Data file for panel b.

Pustelny et al.^21^ ascribed the activating effect of signal peptide removal to changes in overall protein dynamics, which are triggered by the formation of a distinctive hydrogen bond network involving the new N-terminal glutamate residue. To gain insight into the mechanism by which the mutation S154R disables the N-terminal inhibition mechanism of SplB, we solved the crystal structure of SplB S154R and SplB N2K/S154R to a resolution of 1.1 Å and 1.4 Å, respectively (PDB: 6YV5 and 6YV6, Supplementary Table 2). Overall, the structures of the mutant SplB variants are almost identical to those of wildtype pre-SplB (PDB: 4K1S) and activated SplB (PDB: 2VID), previously described by Pustelny et al.^21^ (Fig. 3c), providing no obvious explanation for the observed increase in activity. As a possible measure of differences in the internal protein dynamics, we performed B’-factor comparisons of wildtype and mutant structures but saw no correlation between enzyme activity and B’-factor-based structural flexibility (Suppl. Fig. 3). The structures of SplB mutants S154R and N2K/S154R also lack the extensive hydrogen bond network involving the N-terminus and are thus more similar to the pre-form of SplB (Fig. 3d). Our crystallographic data suggests that SplB S154R and SplB N2K/S154R do not directly mimic the processed form of SplB. Directed evolution using A2M^cap^ has thus provided an outcome that could not have been predicted. If complex dynamic scenarios are responsible for the observed different activities of seemingly identical ground state structures, rational computational approaches would currently be unable to design such subtle differences that directed evolution has identified.

### Mapping of specificity domains of Spl proteases using a SCHEMA shuffling library

A2M bait sequences can be readily modified to accommodate protease recognition sites of interest. Thus, A2M^cap^ should enable directed evolution for new or modified protease specificities by using substrate sequences embedded in A2M.

In analogy to antibodies, consisting of variable regions for antigen recognition embedded in a constant framework, we sought to establish a protease framework that would enable streamlined specificity reprogramming. The Spl protease cluster of *S. aureus* provided a starting point for such a framework: Spl proteases are encoded by a single operon of six paralogues (*splA-F*; 47-95% amino acid identity). Despite their close relationship, they display a remarkably wide range of substrate preferences. SplA, SplB and SplD have been characterised, with VWLY↓S, WELQ↓S and RWLL↓T being their preferred substrates, respectively^20,22,23^. More recently, the last two members of the gene cluster, SplE and SplF, were found to cleave the sequences (V/L)WL(H/Q)↓S and (F/Y)X(L/M)↓S, respectively^24,25^. Given the diversity in substrate preference of these closely related enzymes, we speculate that Spl proteases are particularly amenable to specificity engineering.

First, to find out whether display on yeast changes substrate preferences of the Spl proteases, SplA-F were probed with A2M-VWLY↓S, -WELQ↓S or -RWLL↓T. SplA, SplB and SplD showed activity towards their native substrates, although SplD also showed low levels of activity towards the cognate SplA substrate VWLY↓S. As expected, SplC, despite being displayed, did not show activity against any of the three substrates^18^. SplE and SplF showed less specific cleavage, with activity towards all three substrates to different extents (Fig. 4a, Supplementary Fig. 4). Taken together, our A2M^cap^ assay faithfully recapitulated the previously observed specificities of SplA-F.

**Figure 4:**
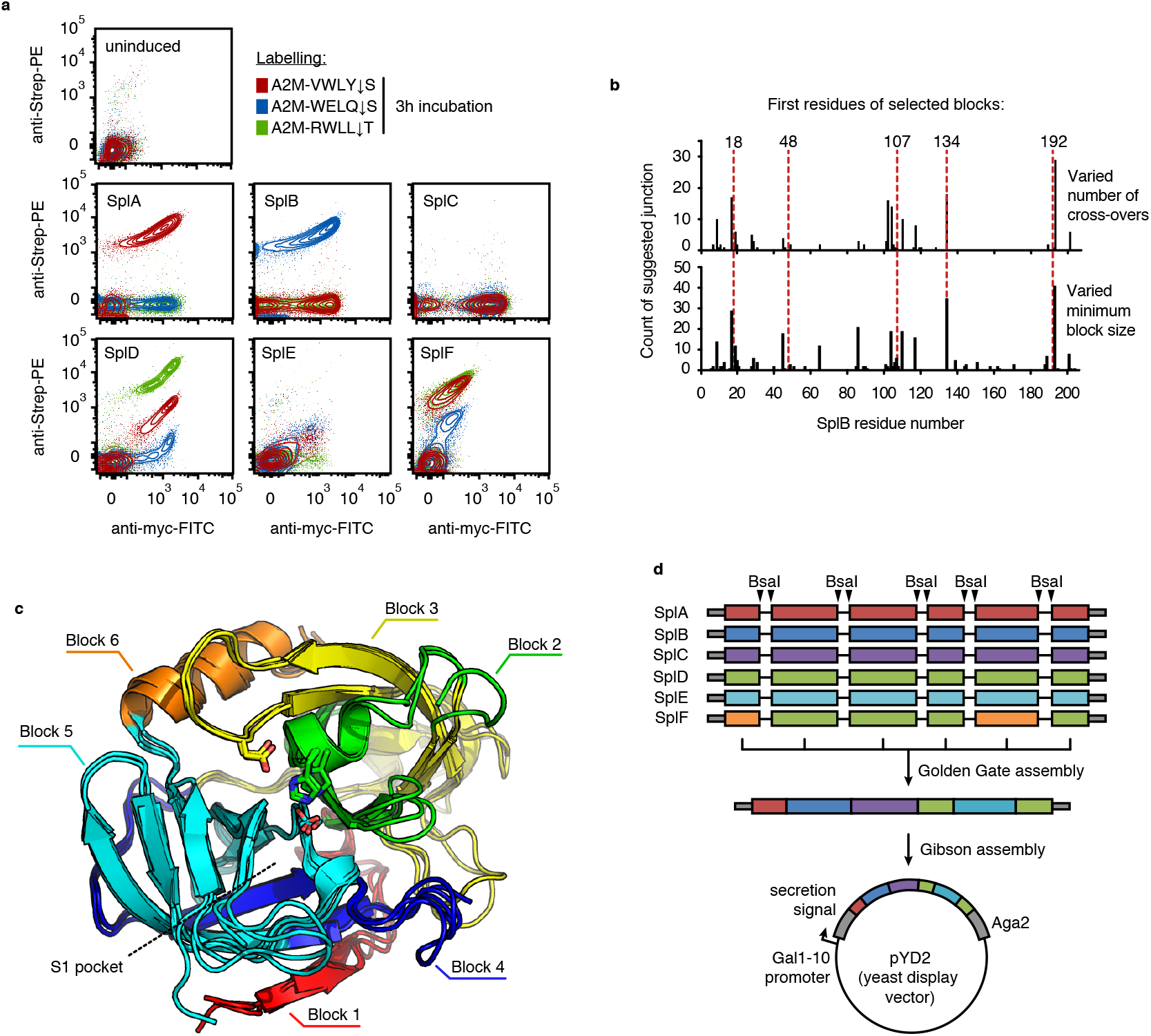
Shuffling library design and assembly for Spl proteases. **(a)** Flow cytometry of Spl protease-displaying *S. cerevisiae* cells probed with substrates A2M-VWLY↓S, -WELQ↓S and -RWLL↓T (representative data of *n*=2 independent inductions and flow cytometry measurements). **(b)** SCHEMA algorithm output for Spl proteases SplA-F run multiple times, varying parameters number of cross-overs or minimum block size. **(c)** Structural alignment of SplA (PDB: 2W7S), SplB (PDB: 2VID) and SplD (PDB: 4INK) with SCHEMA blocks highlighted by color. **(d)** Cloning strategy of SCHEMA library: Each Spl gene was ordered with intervening dual type IIs restriction sites for Golden Gate assembly. The shuffled Spl genes were cloned into the yeast display vector pYD2. FITC = Fluorescein isothiocyanate, Strep-PE = Streptavidin-Phycoerythrin. Source data is provided as a Source Data file for panel 4e.

For Spl proteases to serve as a platform for specificity reprogramming, a better understanding of their substrate recognition and a stable protein framework would be desirable. To address these two aspects, we turned to a domain shuffling approach since this has previously yielded stabilized protein variants^26^ and, moreover, appeared suitable for mapping the specificity determining domains of Spl proteases. We used the SCHEMA algorithm^27^ to identify non-disruptive crossover points, five of which were finally selected based on their SCHEMA energy profile and the level of sequence conservation (Fig. 4b, Supplementary Fig. 5a and b). Examining the suggested crossover points on the structural level revealed that the SCHEMA algorithm had left subdomains largely intact, including the subdomain previously recognised as being important for substrate recognition in Spl proteases (coloured in cyan, Fig. 4c^20,22^).

The Spl shuffling library, with a theoretical diversity of 22500 (= 6×5×5×5×6×5; SplD and SplF have identical amino acids in block 2,3,4 and 6), was generated via Golden Gate cloning and displayed on yeast cells (Fig. 4d, Supplementary Fig. 5c-e). Sequencing of the input library using circular consensus sequencing with Oxford Nanopore technology (Fig. 5a)^28^, revealed the presence of 13,994 different Spl chimeras (62% of the theoretically possible combinations) and an acceptable overall recombination preference (the most overrepresented variant was 18-fold above the median read-count) (Fig. 5b). Flow cytometry of the input library probed with the three cognate substrates A2M-VWLY↓S, -WELQ↓S and -RWLL↓T (previously described for the parent Spl proteases SplA, SplB and SplD, respectively^20,22,23^) showed clones with activities ranging from undetectable to high (Fig. 5c), suggesting high functional diversity of the SCHEMA library.

**Figure 5:**
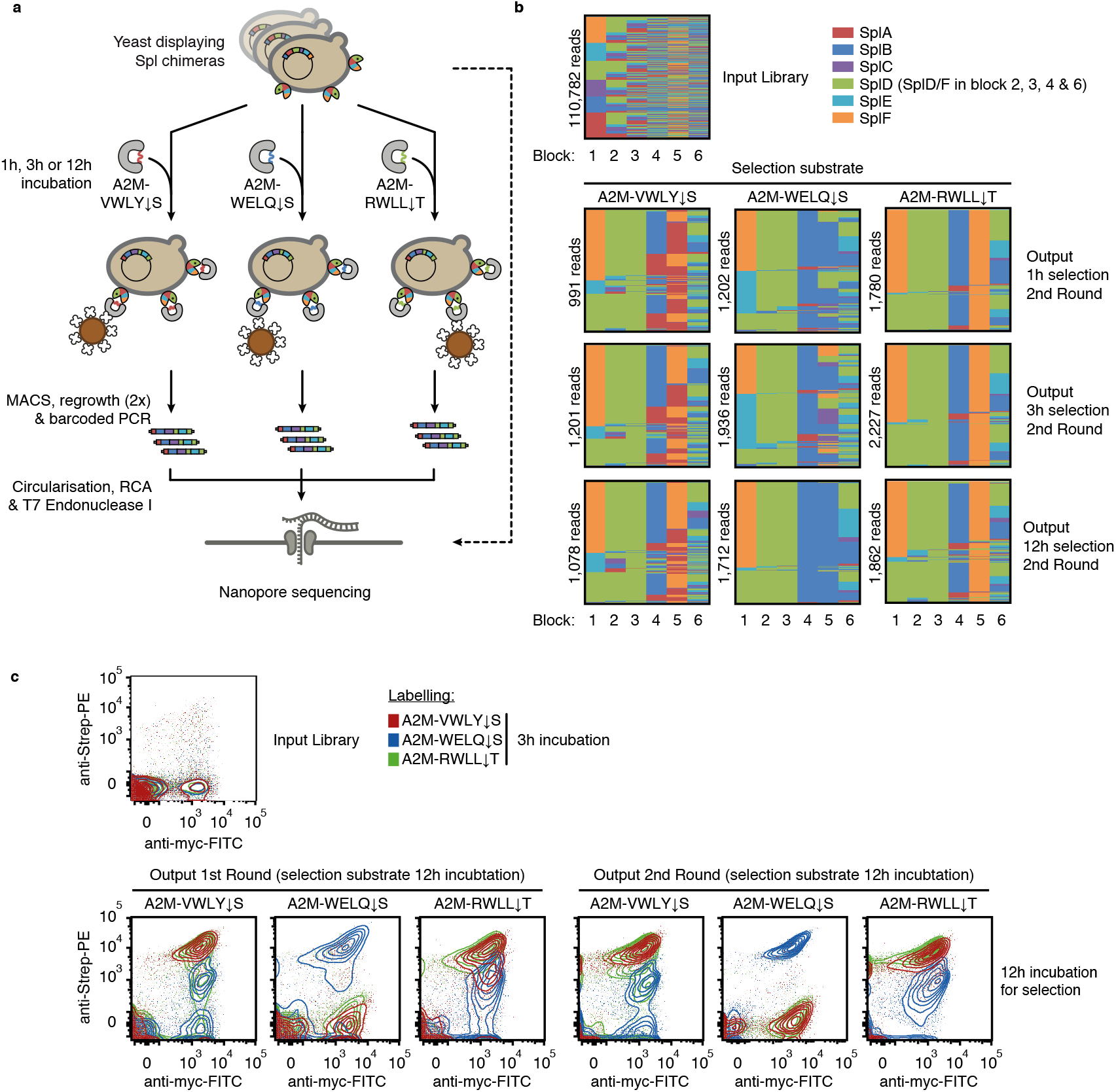
A2M^cap^-based selection of SCHEMA-shuffled Spl proteases. **(a)** Schematic of selection strategy for SCHEMA shuffled proteases using three different substrates and nanopore-based sequencing **(b)** Sequence analysis of input and output populations. Within each graph, sequences of shuffled variants are depicted as lines coloured by block ID at each position. The number of mappable INC-seq reads is shown for each condition/population. **(c)** Flow cytometry of input and output populations probed with the three different A2M substrates used during selection (representative data of *n*=2 independent inductions and flow cytometry measurements for 1^st^ round and input library, n=1 for 2^nd^ round). RCA = rolling circle amplification, PCR = polymerase chain reaction, MACS = magnetic activated cell sorting, FITC = Fluorescein isothiocyanate, Strep-PE = Streptavidin-Phycoerythrin. Source data is provided as a Source Data file for panel b and c.

To identify the subdomains responsible for substrate recognition, the library was subjected to selections using the three substrates A2M-VWLY↓S, -WELQ↓S and -RWLL↓T and three different incubation times (1h, 3h and 12h) (Fig. 5a). After two rounds of MACS, the output populations showed increased activity towards the respective A2M selection substrate (Fig. 5b, Supplementary Fig. 6). Consistently, frequency distributions at each block position shifted compared to the input library. For each substrate, the three different incubation times (1h, 3h and 12h) produced similar enrichment patterns, but shorter incubation resulted in increased background levels. In all selections, we observed an enrichment of SplD/F parent sequences in block 1-3 and SplB parent sequences in block 4, reaching on average a combined ~92% and ~85% frequency, respectively. In block 6, close to equal distributions of SplB, SplE and SplD/F sequences were observed, again, independently of the substrate used for selection (Fig 5c). Thus, a common Spl framework with the block sequence (FDDBX[B/D/E]) was selected with all three different substrates.

In contrast to block positions 1-4 and 6, block 5 showed substrate-specific enrichment corresponding, at least partially, to the described Spl substrate preferences. Specifically, block 5 sequences of SplA or SplF were enriched when selection was performed with the cognate SplA substrate A2M-VWLY↓S (up to 33% and 62%) while SplB sequences were dominant in selections using its cognate substrate A2M-WELQ↓S (up to 97%). Not following this pattern was the selection with A2M-RWLL↓T where SplF block 5 sequences were dominant (up to 98%). The substrate preference for SplF has not been previously characterized but SplD and SplF are the two Spl proteases with highest similarity (94.6% identical, 84.7 % identical in block 5). Sequences of SplC, a parent for which the specificity profile is also not known, were depleted from the output sequences (Fig. 5b and c). Collectively, the enrichment data from the three substrates suggest that block 5 is the predominant specificity-determining block.

### Specificity switching of Spl protease by block 5 domain replacement

To address the role of block 5 for Spl protease specificity, we constructed six chimeric Spl variants with the block sequence FDDB<u>X</u>D (<u>X</u> standing for block 5 sequences of SplA-F). When tested via yeast surface display, the specificity profiles of the chimeras FDDB<u>B</u>D, FDDB<u>D</u>D, FDDB<u>E</u>D and FDDB<u>F</u>D mirrored those of the respective parent protease (Fig. 6a). Thus, the block 5 subdomain appears sufficient for substrate selection for the parent proteases SplB, -D, -E and -F.

**Figure 6:**
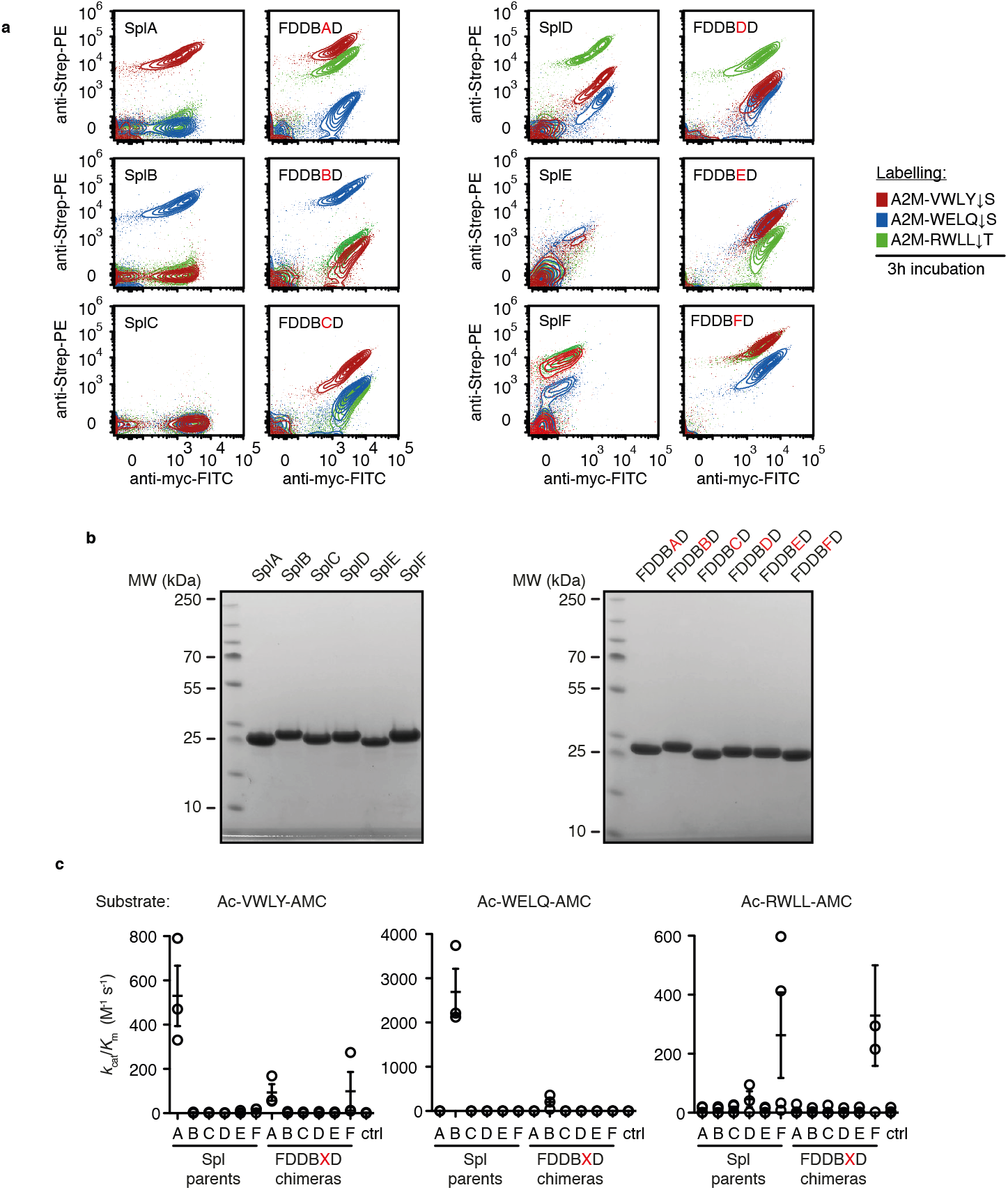
Characterisation of Spl protease chimeras. **(a)** Flow cytometry of parent Spl proteases and chimeras with ID of variable block 5 position indicated in red (representative data of *n*=3 independent inductions and flow cytometry measurements). **(b)** SDS-PAGE of E. coli-purified parent and chimera Spl proteins used for in vitro characterization (native N-termini generated via SUMO-tag removal strategy as described in Methods section Protein expression and purification, result of *n*=1 protein purification). **(c)** Activity measurements (pseudo-first order kinetics) of purified Spl parent proteases and chimeras measured at 0.5-2 μM enzyme and 2.5-5 μM fluorogenic peptide substrates (Ac: acetyl-; AMC: 7-Amino-4-methylcoumarin; *n*=3 independent measurements of the same protein purification, mean with SEM). Source data are provided as a Source Data file for panels a,b,c and e. FITC = fluorescein isothiocyanate, Strep-PE = streptavidin-phycoerythrin, MW = molecular weight, kDa = kilodalton.

FDDB<u>A</u>D and FDDB<u>C</u>D are the two exceptions where the substrate specificity or activity of the FDDB<u>X</u>D chimeras notably differed from the parent proteases. While SplA had shown exquisite specificity for VWLY↓S, FDDB<u>A</u>D also showed activity towards RWLL↓T (Fig. 6a). This suggests that other subdomains apart from block 5 contribute to specificity fine-control in case of SplA. The parent protease SplC so far remains elusive in terms of activity and substrate specificity. For FDDB<u>C</u>D, we found that it has activity towards the cognate SplA-substrate VWLY↓S (Fig. 6a). The SplC block 5 thus appears to be, in principle, substrate binding- and cleavage-competent but non-functional in its native domain context.

We purified the six FDDB<u>X</u>D chimeras and the Spl parent proteases (Fig. 6b) and assessed their *in vitro* activity towards the three test substrates VWLY↓S, WELQ↓S and RWLL↓T. For the parent Spl proteases, the activity profiles were as described for SplA and SplB with *k*_cat_/*K*_m_(VWLY↓S) ~530 ±136 M^-1^s^-1^ and *k*_cat_/*K*_m_(WELQ↓S) ~2690 ±525 M^-1^s^-1^, respectively. Towards their non-cognate substrates, the activity of SplA and SplB was statistically indistinguishable from control reactions without enzyme^21,22^, as was the case for SplC and SplE towards any of the three substrates as expected^18,24^. SplD showed relatively low overall activity but was specific for RWLL↓T (*k*_cat_/*K*_m_ ~45 ±26 M^-1^s^-1^)^23^. In contrast to our A2M^cap^-based results, SplF showed a preference for RWLL↓T (*k*_cat_/*K*_m_ ~263 ±145 M^-1^s^-1^) over VWLY↓S (*k*_cat_/*K*_m_ ~13 ±5.6 M^-1^s^-1^) and its activity towards WELQ↓S was undetectable (Fig. 6c). These results suggest that A2M^cap^ captures the substrate preferences of the parent Spl proteases with some differences for SplD and SplF.

The six FDDB<u>X</u>D proteases showed overall reduced activity but mostly preserved substrate preferences, the latter being expected from the A2M^cap^ flow cytometry data shown in Fig. 6a. Specifically, FDDB<u>A</u>D and FDDB<u>B</u>D were ~5 and ~10-fold less active than their parent proteases towards VWLY↓S (*k*_cat_/*K*_m_ ~93 ±37 M^-1^s^-1^) and WELQ↓S (*k*_cat_/*K*_m_ ~201 ±90 M^-1^s^-1^), respectively. With respect to specificity switch achieved through block 5 exchange, FDDB<u>A</u>D activity for WELQ↓S and, *vice versa*, FDDB<u>B</u>D activity for VWLY↓S was undetectable, as shown above the parent proteases SplA and SplB. Similar to the low *in vitro* activity of SplD, FDDB<u>D</u>D was not detected for the cognate substrate RWLL↓T or the other two test substrates. Likewise, FDDB<u>E</u>D activity could not be detected towards any of the three substrates as expected (Fig. 6c). FDDB<u>F</u>D displayed a similar activity profile as its parent SplF (VWLY↓S: *k*_cat_/*K*_m_ ~99 ±87 M^-1^s^-1^; and RWLL↓T: *k*_cat_/*K*_m_ ~329 ±170 M^-1^s^-1^), which (like SplD) showed activity for RWLL↓T and to a smaller extent for VWLY↓S, both *in vitro* and in our A2M^cap^ data. The low VWLY↓S-directed activity of FDDB<u>C</u>D that was apparent from the A2M^cap^ assay was not detectable *in vitro* (Fig. 6a and c). Taken together, the combined data from the A2M^cap^ assay and the *in vitro* assessment using purified protein confirms that FDDB<u>X</u>D is a functional scaffold allowing specificity switching for Spl proteases. While the A2M^cap^ assay enables highly sensitive profiling of substrate preferences, tests using purified enzyme are necessary to evaluate catalytic activities.

## Discussion

This work establishes A2M^cap^ as a framework for efficient and rapid directed evolution of proteases. Its sensitivity makes A2M^cap^ suitable for identifying weak initial promiscuous activities to kick directed evolution campaigns off. Selection of protein catalysts using turnover-based capture probes using display technologies has previously been explored in context of catalytic antibodies, phosphatases, β-lactamase, nucleases and also proteases^29–32^. For the latter chemical probes or naturally occurring suicide inhibitors of proteases have been described: similar to the way antigens are used non-covalently during binder selections (i.e. for panning or labelling)^10,31^ they serve as covalent capture agents. Covalent capture in A2M^cap^ increases the degrees of experimental freedom, as – in contrast to inhibitors previously employed for protease selections – A2Ms do not directly interfere with catalysis. In fact, protease capture by A2Ms occurs independently of the underlying catalytic mechanism of the protease and only after the rate-limiting step of the reaction (i.e. cleavage of the peptide bond)^11,33^. Thus, A2M are ‘broad spectrum inhibitors’, which capture their protease preys after product formation is complete, making them particularly suitable for the selection of proteases in directed evolution applications. Selection is direct for product formation, rather than a proxy (e.g. noncovalent surface capture^34^, infectivity^5^ or a triggered reaction^30^).

The use of yeast display, in which on the order of 1-10 × 10^4^ enzyme molecules are present on the surface of a cell, sets up a dynamic range (defined by an equivalent number of single turnover reactions that can be followed by the capture of A2M molecules before saturation is reached). Thus activity improvements can be selected, as shown by the order of magnitude activity enhancement for SplB that obviated activating cleavage of its signal peptide. The resulting mutants are useful for situations where the otherwise necessary activation is difficult or impossible to realize, e.g. when an N-terminal fusion partner is required. Nevertheless, for fast enzymes, A2M^cap^ may reach its limits in terms of quantitative discrimination of catalysts in a library. In such cases, assays that measure multiple turnover using fluorogenic substrates in an (ultra)-high-throughput fashion are more suitable^35^. Alternatively, different selection regimes can be realized by varying substrate concentration and incubation time and by using substrate mixtures or multi-specific A2Ms with combined bait sequences.

A2M^cap^ appears particularly suitable for specificity reprogramming given its high sensitivity and, therefore, the ability to detect weak initial activities. Blum et al.^5^ have outlined a strategy for gradual altering of specificity (involving increasingly different substrates that serve as stepping-stones during the directed evolution campaign) that should also be feasible with A2M^cap^. This may involve negative selection using A2M substrates with non-desired bait sequences.

The throughput of A2M^cap^ synergises with the ability to obtain high-throughput sequencing information on a similar scale to gain insight into molecular recognition principles^36^. This was demonstrated here for an Spl SCHEMA library against multiple substrates in parallel, which led to the discovery of specificity determinants of Spl proteases. Since block 5 emerged as the dominant contributor to substrate specificity, a diversification strategy focused on block 5 in the identified permissive protein scaffold may be promising for substrate specificity engineering. Our findings enable a better molecular understanding of Spl proteases and therefore, ultimately, of their role in *S. aureus* virulence and immune-evasive properties^37^. This is exemplified by our unexpected finding that block 5 of SplC mediates the recognition of VWLY↓S as a substrate. Since previous attempts by Popowicz et al.^18^ to identify the natural substrate of the parent protease SplC were unsuccessful, the activity of FDDB<u>C</u>D might serve as a useful starting point for future studies on SplC (Fig. 4d and e). We envision that specificity engineered Spl proteases would be particularly useful in synthetic biology applications, e.g. for proteolysis-based logic^38–40^.

Since the capture mechanism of A2Ms is independent of the catalytic mechanism of the cleaving protease, A2M^cap^ is, in principle, applicable to most protease classes. Nevertheless, there may be cases where capture is inefficient or absent. Apart from bacterial A2Ms used here, bait-engineered human A2Ms have also been described recently and a diverse set of proteases showed activity towards these substrates. Albeit their purification being more cumbersome, these human A2Ms may be an alternative option should capture efficiencies be an issue with bacterial A2Ms^41^.

While demonstrated in combination with yeast surface display, A2M^cap^ is platform-independent and compatible with other commonly used protein display formats. For example, *in vitro* display formats (ribosome and puromycin display, SNAP display^42^) may be particularly interesting in the context of proteases to circumvent host toxicity and thus loss of promiscuous intermediates in directed evolution campaigns^43,44^. Taken together, A2M^cap^ will open new avenues for the development of proteases as biological precision tools with reduced off-target activity, higher stability and greater resistance to endogenous inhibitors.

## Methods

### DNA constructs

Sequence files of DNA constructs used in this study can be found online at Benchling [https://benchling.com/philipp_knyphausen/f_/M2tyfz1K-a2m_deposit/]. A table with a description for each construct is provided in Supplementary Table 3. Gene synthesis was carried out at Thermo Geneart. Oligonucleotides and primers were synthesised at Sigma Aldrich. The *yfhM* gene coding for A2M was amplified from *E. coli* BL21 cells.

### Protein expression and purification

A2M constructs were expressed in BirA co-expressing *E. coli* BL21 (DE3) cells at 1-2 L scale (typical yield ~4-8 mg/L). Cells were grown at 37 °C until an OD_600_ of 0.6-0.8, at which point 250 μM IPTG and 40 mg/L biotin (in DMSO) were added and the temperature was reduced to 20 °C. Cells were harvested after 16 h, resuspended in PBS/10 mM imidazole and lysed by sonication. After removal of the cell debris by centrifugation for 30 min at 12.000 × *g*, the lysate was passed over ~1 ml of Ni-NTA resin, washed with 20 ml of PBS/20 mM imidazole/500 mM NaCl and eluted in 2.5 ml PBS/250 mM imidazole. A buffer exchange into PBS was performed with a PD-10 column, the protein was concentrated with a 100 MWCO centricon and finally aliquoted, snap-frozen in liquid nitrogen and stored at −80 °C.

Purification of Tamavidin-2-HOT^45^ was carried out as described for A2M substrates but in standard *E. coli* BL21 (DE3) cells and without added biotin. Likewise, His_6_-tagged SplB variants (from the epPCR directed evolution campaign, cloned into pRSF-NheI) and SUMO-tagged Spl constructs were expressed in *E. coli* BL21 (DE3) cells, which were however induced with 1 mM IPTG and grown at 20 °C for 3-4 h before cell collection. For the generation of native N-termini, N-terminally SUMO-tagged and C-terminally His_6_-tagged Spl constructs in pHAT5^46^ were co-expressed with GST-Ulp1 (in pRSF-Duet-1). Growth and purification were performed as for the SplB variants described above.

### Yeast display, labelling and selections

*S. cerevisiae* EBY100 cultures were grown and induced as described^47^. Glucose cultures were grown for ~16 h at 30 °C and passaged at a 1:10 dilution into SG-CAA medium for induction followed by incubation for 2 days at 20 °C. Culture volumes and sample sizes were adjusted to ensure an at least ten-fold excess of cells relative to the library diversity (OD_600_ 1 ~ 3 × 10^7^ cells/ml). Media were supplemented with alternating antibiotics after MACS to reduce the risk of bacterial contamination (50 μg/ml Kanamycin, 25 μg/ml chloramphenicol or 1x Pen/Strep, Gibco). Transformation was carried out by electroporation for libraries^48^ and using the Frozen-EZ Yeast Transformation II Kit (Zymo) for individual constructs.

### Tamavidin coated magnetic beads used for MACS

Tamavidin-2-HOT was covalently coupled to paramagnetic carboxy beads (Ø 1 µm; PS-MAG-COOH-S2472, microParticles, Berlin). Beads (100 mg) were washed with water (with 0.02% v/v Tween-20), then resuspended in 1 mL water (with 0.02% v/v Tween-20). To the bead suspension was added 0.5 mL of 750 mM of N-(3-dimethylaminopropyl)-N’-ethylcarbodiimide hydrochloride (EDC) in water (Sigma-Aldrich) and the mixture was incubated for 20 minutes. The supernatant was removed, the beads were washed once with water (with 0.02% Tween-20), before resuspension in 5 mL of 25 mM sodium phosphate buffer (pH 6; with 0.02% Tween-20). Subsequently, Tamavidin-2-HOT protein (1.5 mL of 308 µM (tetramer concentration) in PBS) was added and the tube was left on a roller at room temperature for four hours. Finally, the beads were washed with 0.5 M Tris-HCl (pH 8, with 0.05% Tween-20) and incubated in this buffer for 10 minutes in, followed by washing with PBS (with 0.05% Tween-20).

### Magnetic activated cell sorting (MACS)

For MACS of the epPCR library, cells from 50 ml of the induced library (~1 × 10^9^ cells) were washed once with 50 ml sterile PBS/0.5% BSA and then labelled in 1 ml 2.5 μM A2M-WELQ↓S over night on roller at room temperature. Labelled cells washed three times with 12 ml sterile PBS/0.5% BSA and finally resuspended in 5 ml PBS/0.5% BSA. An input sample was taken for later analysis. Then, ~1 × 10^7^ Tamavidin-coated beads were added, followed by incubation for 1.5 h at room temperature on a roller. Beads were collected on a magnet, transferred to a 1.5 ml tube and washed five times. As the last step, beads were resuspended in 50 μl PBS/0.5% BSA and transferred into 1 ml SD-CAA with Pen/Strep for recovery of the cells (16 h at 30 °C). The second round and the shuffling library selections were carried out similarly but with cell/bead numbers scaled-down (~1 × 10^7^ cells, ~1 × 10^6^ beads).

### Enzyme assays & kinetics

Michaelis-Menten kinetics of pre-SplB mutants were measured with the peptide substrate Ac-WELQ-AMC (Ac: acetyl-; AMC: 7-Amino-4-methylcoumarin, stock concentration: 26 mM in DMSO, concentration range: 13-1161 μM, Genscript) at an enzyme concentration of 125 nM to 2.5 μM using a Tecan infinite 200Pro (excitation wavelength 339 nm, emission wavelength 439 nm). Before curve fitting in Graphpad Prism, fluorescence signals were background-subtracted from a control time course without enzyme and the corresponding substrate concentration. In addition, the Ac-WELQ-AMC substrate showed a non-linear signal increase with increasing concentration, thus requiring correction with a calibration curve derived from the fluorescence values after complete turnover of the substrate.

### Error-prone PCR library generation

Random mutagenesis was carried out in a 50 µl reaction with the JBS dNTP-Mutagenesis Kit (Jena Bioscience) with 5 µM dNTP analogs, 500 µM natural dNTPs, 100 ng pCT-SplB template and 0.5 µM primers PK417+PK418. The PCR reaction was run with the following settings: (1) 3 min at 92 °C, (2) 1 min at 92 °C, (3) 1.5 min at 55 °C, (4) 5 min at 72 °C, (5) 4x to step 2 and (6) 10 min at 72 °C. After over-night treatment with 2 µl DpnI at 37 °C, 20 µl of the epPCR served as a template for a second 1 ml standard PCR with 25 cycles using BioTaq polymerase (Bioline) and primers CW1&2 as previously described^49^. Digested vector was prepared from pCTcon2 bearing an insert with a unique SphI site. After over-night digestion of 50 µg of plasmid DNA with 9 µl SphI in 300 µl, 3 µl of BamHI and NheI were added and digestion was continued for 12 h. Another 3 µl of BamHI and NheI were added for a final over-night incubation. The PCR product and digested vector were SPRI purified, eluted in 100 µl ddH_2_O and dried in a speedvac. The DNA resuspended in 20 µl ddH_2_O (~1-3 µg/µl) was ready for yeast transformation by electroporation. Before yeast transformation the same DNA material was used for Gibson assembly (carried out as described by Gibson et al.^50^) and subsequent *E. coli* transformation. Twelve single colonies were picked for Sanger sequencing, which showed an average error rate of 1.9 mutations/gene with a range from 0-4.

### SCHEMA shuffling library generation

SCHEMA tools were used as described^51^. The algorithm was run with a multiple sequence alignment of SplA-F and the SplB structure (PDB 2VID) as input and varying settings for number of crossovers and minimum number of mutations per block. For generation of the SCHEMA shuffling library, *splA-F* genes containing tandem-BsaI restriction sites at cross-over points were ordered as synthetic Gene Strings (GeneArt) with Gibson overhangs for cloning into PstI/NheI digested pYD2 vector. Two mutations had to be introduced into SplA (K48S/N49D) and one into SplE (A48D) to facilitate cloning of the library (Supplementary Fig. 5c and e). These mutations were also included when the parent variants were tested initially (Fig. 4a). Each variant was amplified with primers PK421&422 using Q5 High-Fidelity 2X Master Mix (NEB) in a 200 µl reaction followed by DpnI digestion for 2 h. PCR products were SPRI purified, eluted in 50 µl and quantified using a Nanodrop spectrometer. The Golden Gate reaction was carried out in a 100 µl volume with 1 µg of each *spl* PCR product, 10 µl T4 DNA ligase buffer (10x), 5 µl T4 DNA ligase (both NEB) and 5 µl FastDigest Eco31I (BsaI isoschizomer from ThermoFisher). The reaction incubated in a thermocycler for 12 h with 10 min steps at 37 °C and 16 °C, followed by a final heat inactivation step at 80 °C for 10 min. The reaction was SPRI purified and eluted with 50 µl of an Eco31I containing digestion mix (2 µl enzyme, 5 µl 10x FastDigest Buffer, 43 µl ddH_2_O). This reaction was run for 3 h at 37 °C and subsequently loaded onto a 1 % agarose gel. The band at ~770 bp was extracted from the gel and eluted in 20 µl elution buffer. The final concentration was 15 ng/µl. Vector was prepared from 10 µg pYD2-SplB-S157A (bearing a unique NsiI site) by digesting with PstI, BamHI and NsiI and SPRI purification. An 80 µl Gibson assembly with 250 ng of the prepared insert and 650 ng of digested vector (molar ratio ~3:1) was run for 1 h at 50 °C, SPRI purified and eluted in 10 µl. Four 25 µl aliquots of E. cloni 10G ELITE Electrocompetent Cells (Lucigen) were transformed with 2.5 µl of the eluate each. This transformation gave rise to ~3.4 × 10^6^ cfu (~150 × oversampling of theoretical diversity). DNA from single colonies was used to verify successful shuffling. Colonies were scraped and plasmid DNA was prepared from a 50 ml culture of the pool. Since circular plasmid DNA is less efficient at transforming yeast^52^, 20 µg were linearised with XhoI/SacI (as above for pCTcon2 but scaled down accordingly) and SPRI purified. An insert downstream of *aga2* was amplified from pYD2 using primers PK463+PK464 in a 1 ml PCR reaction (Q5 High-Fidelity 2X Master Mix, NEB), DpnI digested for 2 h and SPRI purified. The forward primer introduces an N_10_-barcode and an additional PstI site for circularisation. Vector and insert were dried with a speedvac and resuspended in ddH_2_O. The yeast transformation yielded ~ 4.3 × 10^7^ cfu (~1900 × oversampling).

### Flow cytometry

For flow cytometry, ~1 × 10^6^ induced cells were washed with 200 µl PBS/0.5% BSA (30 sec spin in a benchtop minicentrifuge, ~1000 × g) and incubated with 2.5 µM A2M substrate in a 50 µl volume at room temperature on a roller. Cells were then washed three times as was the case for the subsequent labelling steps. Streptavidin-PE (1:200, Biolegend, cat. no. 405204) was first applied for 1 h followed by staining with PE anti-streptavidin (1:200, Biolegend, cat. no. 410503, clone 3A20.2) and anti-c-myc-FITC (1:50, Miltenyi Biotec, cat. no. 130-116-485, clone SH1-26E7.1.3) also for 1 h. Flow cytometry was carried out on a Becton Dickinson FACScan Cytek DxP machine with a 488 nm laser / 530/30 nm filter for FITC and a 561 nm laser / 590/20 nm filter for PE. Flow cytometry data was analysed with FlowJo v10.1.

### DNA preparation, Nanopore high-throughput sequencing

DNA libraries for Nanopore sequencing were prepared by extracting plasmid DNA from ~1 × 10^9^ cells (input library, 30 ml culture) or ~3 × 10^7^ cells (output populations, 1 ml culture) using Zymoprep Yeast Plasmid Miniprep (Zymo). One freeze-thaw cycle (at −80°C) was performed before and one after addition of zymolase (15 µl/ml, 1 h at 37 °C treatment). The input library DNA was purified over four miniprep columns. After elution with 30 µl elution buffer, 5 µl of the plasmid DNA served as template for a 50 µl PCR (Q5 High-Fidelity 2X Master Mix, NEB) with primers PK412+PK421 (input library) or PK412+barcode primer (PK488–PK496 and PK524). Barcode sequences were derived from the PCR Barcoding Kit 96 (Oxford Nanopore). Amplicons were individually gel extracted, eluted in 15 µl and the concentration was determined on a Qubit dsDNA HS Assay Kit (ThermoFisher). To increase sequencing accuracy we employed INC-seq^9^. For INC-seq, amplicons of the selection outputs were pooled (80 ng each, 720 ng total) and PstI-digested in a 50 µl volume for 2 h at 37 °C. Similarly, 1 µg of the input library amplicon was used for the digest. After SPRI purification and elution in 20 µl, the DNA concentration was again determined using via Qubit. Ligation was carried out in a 500 µl volume with 20 µl T4 DNA ligase and 0.5 ng/µl of digested PCR products. After SPRI purification, elution in 20 µl and Qubit measurement (yield ~8 ng/µl), the rolling circle amplification was set up as follows (separately for input and output samples): 20 µl random hexamers (1 mM), 40 µl dNTPs (10 mM), 8 µl template, 40 µl 10X phi29 DNA Polymerase Reaction Buffer (NEB) and 40 µl ddH2O. The reaction was heated to 95°C and cooled to 4°C, supplemented with 16 µl DTT (100 mM), BSA (10mg/ml), 20 µl phi29 DNA Polymerase (10 U/µl, NEB) and incubated for 12 h at 30 °C. The RCA product was SPRI purified (incubation time for 15 min on roller for binding and elution), eluted in 100 µl T7 endonuclease I reaction mix (5 µl T7 endonuclease I, 10 µl NEBuffer 2) and incubated for 1.5 h at 37 °C with gentle rocking (still in presence of beads). After separation on a 1 % agarose gel for 1 h at 60 V, DNA fragments above 10 kb were gel extracted and eluted in 60 µl ddH_2_O. The DNA concentration was determined by Qubit and Nanodrop (yield ~1.5 µg). A repair reaction was run using the PreCR kit (NEB) as described by the manufacturer followed by SPRI bead purification and elution in 51 µl ddH_2_O. Subsequent steps for the preparation of the MinION sequencing run were carried out based on the LSK-108 ligation protocol (Oxford Nanopore) but with prolonged times (15 min) for binding to and elution from AmpureXP beads to improve DNA recovery.

### Sequencing data analysis

Fastq-files were generated using the high accuracy (HAC) base calling setting in MinKNOW and were filtered with NanoFilt 1.9.2 [www.github.com/wdecoster/nanofilt] based on a quality score of >=9 and 3000 bp length. Input library sequencing data obtained from three different runs were combined. After conversion into fasta-format using seqkit 0.12.0 [www.github.com/shenwei356/seqkit], the INC-seq [www.github.com/CSB5/INC-Seq] algorithm was run with the minimum copy number set to three and with the *aga2* sequence serving as anchor sequence^28^. A block-wise BLAST was performed (set to ‘short’ for blocks 1, 2, 4 and 6) against separate BLAST databases of *splA-F* sequences of every block position. The tabular BLAST output (containing the best match per sequence) was filtered for an alignment length of >90% for each block. The block IDs were extracted and combined to generate six-number codes for each sequence (i.e. 222222 for a variant consisting of *splB* at every block position). For de-multiplexing of experiments, INC-seq output files were blasted against a database of barcode sequences (Supplementary Table 4) and the resulting experiment specific sequence IDs were matched to the corresponding block sequences. Sorted block IDs for both the input population and the de-multiplexed output populations were then used to create heatmaps (Seaborn 0.9.0, Python 2.7.16) and to calculate block frequencies (Pandas 0.24.2). For details of the commands and scripts used for this analysis see Analysis_HAC.txt file in Source Data. We detected approximately 3% of vector background sequences consisting of (unshuffled) inactive SplB S157A. About 5% of the sequences still contained at least one BsaI restriction site, suggesting incomplete digestion or re-ligation in the Golden Gate cloning process.

### Crystallisation, data collection and structure solution

Crystallization of SplB N3Q/S154R and SplB N2K/N3Q/S154R was essentially performed as described^21^. Crystals were obtained in PEG3350 30%, Tris-Cl pH 8.0 and flash frozen in liquid nitrogen without cryoprotectant. Diffraction data was recorded in a Diamond Light Source Beamline I04 synchrotron radiation source. The data were processed with the automated data-processing pipeline using XIA2-DIALS^53^.The structure was solved by molecular replacement with Phaser (CCP4 suite 7.1.015)^54^ using a previous structure 2VID. The structure was refined using REFMAC5 (CCP4 suite 7.1.015)^54^ with automatic rebuilding using Buccaneer (CCP4 suite 7.1.015)^55^ and manual editing in Coot (CCP4 suite 7.1.015)^56^. Ligand constraints were generated using Grade^57^. Data-collection and refinement statics are given in Supplementary Table 2. B-factors analysis was performed based on the normalised B-factor (B’-factor) obtained from the BANΔIT server [https://bandit.uni-mainz.de/]^58^.

## Acknowledgements

This work was supported by the EPSRC (EP/H046593/1) and the European Union’s Horizon 2020 research and innovation programme. MRP thanks the Royal Society for a Newton Fellowship. FH is an ERC Advanced Investigator (695669). We thank Diamond Light Source for access to beamline I04 (proposals 18548 and 25402), data from which contributed to this manuscript. We are grateful for the X-ray crystallographic facility at the Department of Biochemistry for access to instrumentation and support. We thank Grzegorz Dubin for providing substrates, plasmids, purified enzyme and guidance for the *in vitro* testing of Spl porteases.

## Author Contributions

PK, MR, LJ and FH designed research. PK and MR performed experiments. PB and MH analysed crystallography data. PK and FH wrote paper with input from all authors.

